# A rapid and automated sample-to-result *Candida auris* real-time PCR assay for high-throughput testing of surveillance samples with BD MAX™ open system

**DOI:** 10.1101/608190

**Authors:** L. Leach, A. Russell, Y. Zhu, S. Chaturvedi, V. Chaturvedi

## Abstract

The multidrug-resistant yeast pathogen *Candida auris* continues to cause outbreaks and clusters of clinical cases worldwide. Previously, we developed a real-time PCR assay for the detection of *C. auris* from surveillance samples (Leach et al. JCM. 2018: 56, e01223-17). The assay played a crucial role in the ongoing investigations of *C. auris* outbreak in New York City. To ease the implementation of the assay in other laboratories, we developed an automated sample-to-result real-time *C. auris* PCR assay using BD MAX™ open system. We optimized sample extraction at three different temperatures and four incubation periods. Sensitivity was determined using eight pools of patient samples, and specificity was calculated using four clades of *C. auris*, and closely and distantly related yeasts. Three independent extractions and testing of two patient sample pools in the quadruplicate yielded assay precision. BD MAX™ optimum assay conditions were: DNA extraction at 75°C for 20 min, and the use of PerfeCTa Multiplex qPCR ToughMix. The limit of detection (LOD) of the assay was one *C. auris* CFU/PCR reaction. We detected all four clades of *C. auris* without cross-reactivity to other yeasts. Of the 110 patient surveillance samples tested, 50 were positive for *C. auris* using the BD MAX™ System with 96% clinical sensitivity and 94% accuracy compared to the manual assay. BD MAX™ assay allows high-throughput *C. auris* screening of 180 surveillance samples in a 12-hour workday.

## INTRODUCTION

*Candida auris*, an emerging multidrug-resistant yeast pathogen, continues to cause outbreaks and clusters of clinical cases worldwide (1, 2). There are ongoing efforts to devise better diagnostic approaches for the rapid detection of *C. auris* in clinical and surveillance samples (3–5). Previously, we developed and validated a manual real-time PCR assay for the direct detection of *C. auris* from surveillance samples at the New York State Department of Health (NYSDOH) Mycology Laboratory (3). The laboratory-developed test (LDT) enabled NYSDOH laboratory scientists and epidemiologists to carry out unprecedented surveillance and testing in hospitals and healthcare facilities in New York City. To date, over 13,000 clinical samples from 151 healthcare facilities and over 1,000 *C. auris* isolates were processed (2). *Candida auris* LDT, standard operating procedures, and validation results were shared extensively with the hospital, commercial, and public health laboratories (S. Chaturvedi, unpublished data). However, the *C. auris* LDT is not amenable to automation and high-throughput screening, and adoption of the LDT has progressed slowly while the affected facilities and sample numbers continue to grow. Therefore, we developed and validated a real-time PCR assay for *C. auris* using BD MAX™ System, a fully integrated and automated platform amenable to direct detection of other fungal pathogens (6, 7).

## MATERIALS AND METHODS

As we aimed to migrate the manual *C. auris* real-time PCR assay to BD MAX™, the primers, probe, and PerfeCTa Multiplex qPCR ToughMix (Quanta Biosciences), were evaluated with the BD MAX™ DNA MMK SPC master mix (3). We optimized the BD MAX™ ExK™ DNA 1 (Plasma/Serum/Urine) DNA Extraction Kit on the BD MAX™ using three different temperatures and four incubation periods. We carried out the entire sample-to-result procedure in the BD MAX™ PCR Cartridges. BD MAX™ assay sensitivity was determined using eight pools of patient surveillance samples (axilla, groin, axilla-groin, and nares swabs) in parallel testing with the manual real-time PCR assay. Pooled samples were used to ensure sufficient volume to test identical samples in triplicate using the manual and BD MAX™ assays. High, medium, and low Ct pools were created by combining equal volumes from individual swab samples with similar Ct values based on the manual assay. All pooled samples were then re-run with the manual assay and the BD MAX™ assay to be consistent with the results. We determined assay specificity by the analysis of four clades of *C. auris* and closely and distantly related yeasts at high and low concentrations. The primers and probe set were previously screened against a more extensive panel comprising closely and distantly related fungi, bacteria, viruses, and parasites (3). The assay precision was determined by testing samples in three independent extractions and testing two patient pools in quadruplicate. Following validation, we used the assay for the screening of 110 individual patient surveillance samples. GraphPad Prism 8 software (GraphPad Software, Inc., La Jolla, CA) was used for statistical analysis. We used student t-test for analysis of the means and considered a *P* value of <0.05 statistically significant. The manual *C. auris* real-time PCR assay was selected as the “gold standard” to assess the diagnostic performance of BD MAX™ assay for the surveillance samples. BD MAX™ assay hands-on time estimation involved the sum of the time required to complete sample preparation, device loading and cleaning, and result review and reporting.

## RESULTS

The initial experiments involved optimization of DNA extraction and amplification steps with the BD MAX™ system. This included initial DNA extraction at three different temperatures, 70°C, 75°C, and 80°C, five different incubation times ranging from 10 to 30 minutes, and amplification using two master mixes, the PerfeCTa Multiplex qPCR ToughMix (manual real-time PCR assay) and BD MAX™ DNA MMK SPC master mix. The ToughMix appeared better than the BD MAX™ DNA MMK SPC master mix as it allowed more efficient amplification of *C. auris* DNA (Supplementary Figure 1A). Next, we fine-tuned DNA extraction time and temperature using the ToughMix. Our results revealed that the optimum sample DNA extraction condition was 75°C for 20 min on the BD MAX™ (Supplementary Figure 1B). The selected extraction condition was further evaluated using two pooled surveillance samples (axilla, groin, axilla-groin, and nares) with high and medium Ct values in quadruplicate and results showed excellent reproducibility (Supplementary Table 1). In expanded testing, BD MAX™ assay was highly sensitive with the limit of detection (LOD) of one *C. auris* CFU/PCR reaction (Figure 1). The assay detected all four known clades of *C. auris* with similar Ct values for the same concentration of cells (Supplementary Table 2). The assay was highly specific as no cross-reactivity was seen against two closely related yeasts, *C. haemulonii* and *C. duobushaemulonii*, and 2 distantly related yeasts, *C. albicans* and *C. glabrata*, tested at high (1 ×10^4^ cells/PCR reaction) and low (1 × 10^1^ cells/PCR reaction) concentrations (Supplementary Table 2). The assay was highly reproducible as it produced consistent Ct values on three different days of testing (Supplementary Table 3A), and within the same day of testing (Supplementary Table 3B). Upon completion of the test validation, we tested 110 individual patient surveillance samples; 50 were positive for *C. auris* by BD MAX™ System, resulting in 96% clinical sensitivity and 94% accuracy when compared to the manual real-time PCR assay (Table 1). The Ct value distributions of 110 individual patient surveillance samples showed excellent correlation between two methods (Figure 2). We estimated the automated sample-to-result *C. auris* BD MAX™ assay to allow the screening of 180 samples in a 12-hour workday.

**Figure 1.**
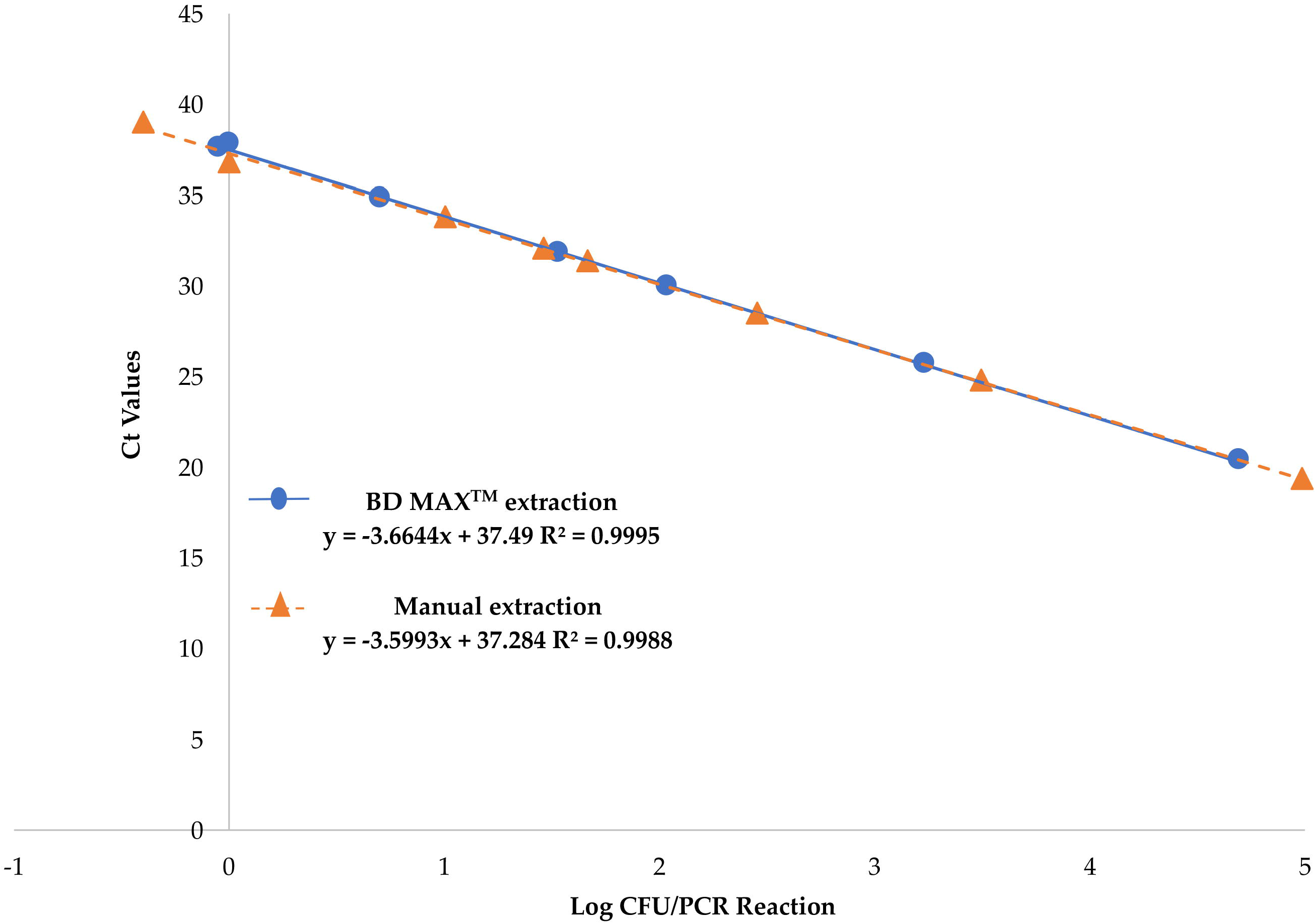
BD MAX™ *Candida auris* assay sensitivity. The pool of patient surveillance samples ranging in *C. auris* CFU from 5 × 10^−1^ - 5 × 10^5^/PCR reactions were run in duplicate on three different days. The assay was linear over six orders of magnitude, and the limit of detection limit was one CFU/PCR reaction on BD MAX™ and manual assay.

**Table 1.** Comparison of Test Performance of Individual Patient Surveillance Sample

| BD MAX™ Real-Time PCR Result | Manual Real-Time PCR Result |  | Accuracy (%) | Sensitivity (%) | Specificity (%) | PPV (%) | NPV (%) |
| --- | --- | --- | --- | --- | --- | --- | --- |
|  | Positive | Negative |  |  |  |  |  |
| Positive | 45 | 5 | 94% | 96% | 92% | 92% | 97% |
| Negative | 2 | 58 |  |  |  |  |  |

**Figure 2.**
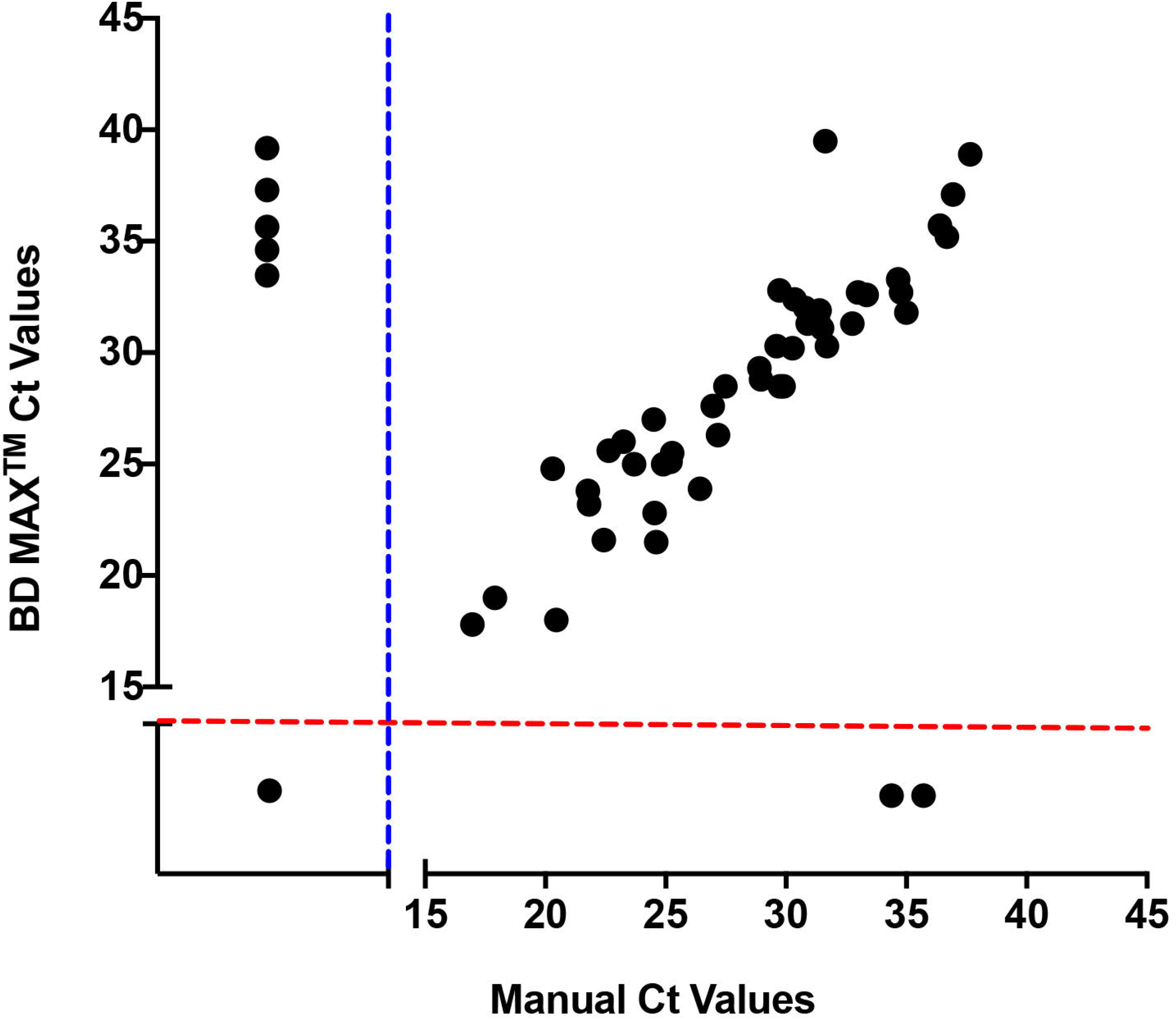
Side-by-side comparison between *Candida auris* BD MAX™ real-time PCR and manual real-time PCR assays. Ct distribution of 110 individual patient surveillance samples showed that BD MAX™ real-time PCR assay had an excellent correlation for the detection of *C. auris* DNA with manual real-time PCR assay for most of the samples (r = 0.9955).

## DISCUSSION

We developed and validated the first automated sample-to-result real-time PCR assay for high-throughput testing of *C. auris*. The highlights of this study are: easy migration of the manual real-time PCR assay to an automated platform, the consistent performance of primers, probe, and master mix from our manual assay, comparable accuracy, sensitivity and specificity, and easy integration in the laboratory workflow. These findings also confirm the suitability of BD MAX™ platform for direct detection of fungal pathogens from clinical specimens as reported earlier for *Pneumocystis jirovecii* and *Coccidioides immitis* (6, 7). There was indication that BD MAX™ performed better than the manual assay for a few samples with high Ct values possibly due to the elimination of many hands-on steps and the use of purified DNA. Although the samples tested in the present study belonged to *C. auris* South Asia clade I, our primers and probe have performed equally well with other well-known *C. auris* clades in the validation panel and the manual assay described earlier (3). Ahmad et al. (2019) partially modified our manual real-time PCR assay by carrying out DNA extraction from suspected *C. auris* surveillance samples on MagNA Pure 96 automated extraction system (Roche Diagnostic System, Indianapolis, IN, USA); these authors reported a good correlation of the modified test findings with the results obtained by cell culture and MALDI-TOF-MS identification (4). The authors further claimed that partial automation of the real-time PCR assay allowed testing of approximately 200 samples per day. However, the trend in infectious disease diagnosis is towards full automation, multiplexing, and miniaturization of assay systems to facilitate rapid point-of-care (POC) diagnosis (8, 9). It is encouraging to note that another group has successfully migrated manual real-time PCR assays to multiple sample-to-result platforms such as BD MAX™, ELITe InGenius (ELITechGroup), and ARIES (Luminex) simultaneously(10). In summary, the development and validation of the rapid and automated sample-to-result *C. auris* BD MAX™ open system assay would allow for the wider adaptability and availability of surveillance testing at the front-line laboratories.

## ACKNOWLEDGMENTS

This work was supported by the Centers for Disease Control and Prevention-Antibiotic Resistance Lab Network grant (NU50CK000423) and the Clinical Laboratory Reference System, Wadsworth Center, New York State Department of Health.

L. Leach assisted in the assay’s design, performed the experiments, and tabulated and analyzed the data. A. Russell assisted in the assay’s design, performed the experiments, and participated in the analysis of data, and edited the manuscript. Y. Zhu performed experiments and tabulated and analyzed the data and edited the manuscript. S. Chaturvedi conceived and designed the study, interpreted data, and edited the manuscript. V. Chaturvedi conceived and designed the study, interpreted data and wrote the manuscript. All authors commented upon the final manuscript.

