## Supplementary material for "A rapid and automated sample-to-result *Candida auris* real-time PCR assay for high-throughput testing of surveillance samples with BD MAX™ open system"

**Supplementary Figures.**

Supplementary Figure 1A. Comparison of efficacy of two different master mixes on the BD MAX™️ platform. The PerfeCTa Multiplex qPCR ToughMix used in manual real-time PCR assay was compared with BD MAX™️ DNA MMK SPC master mix with DNA extracted at 70°C and 80°C. The limited test indicated the ToughMix was better than BD MAX™️ DNA MMK SPC master mix as it allowed more efficient amplification of *C. auris* DNA for samples with high Ct values.

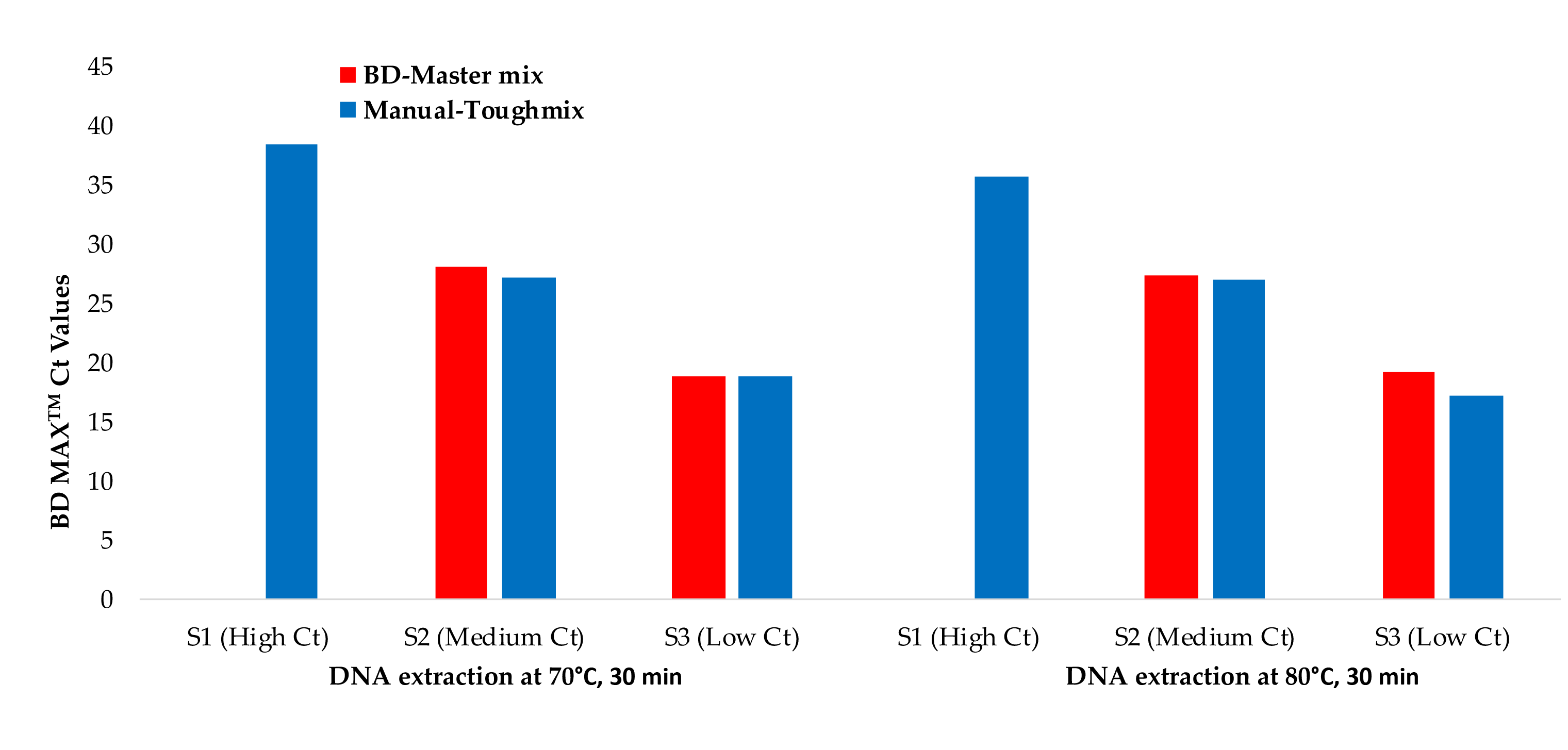

Supplementary Figure 1B. Further Performance of BD MAX™️ with PerfeCTa Multiplex qPCR ToughMix. Two patient pools (high and medium Ct counts) were tested at three different temperatures and four incubation periods. The results revealed that optimum sample DNA extraction condition was 75°C for 20 min on the BD MAX™, highlighted with the green box.

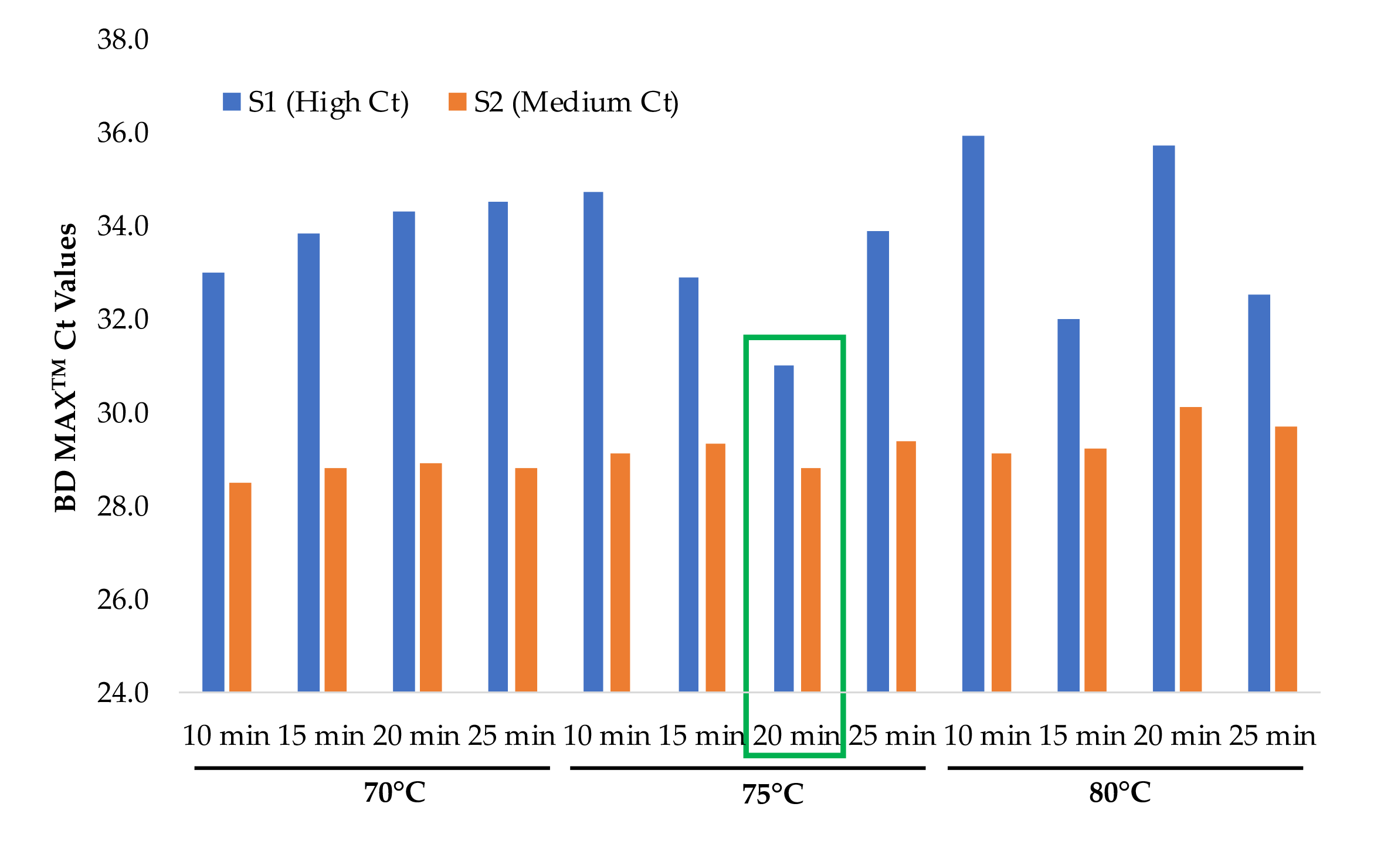

**Supplementary Table 1. BD MAX^™^ Assay reproducibility with ToughMix at 75°C for 20 min.** Two pools of surveillance samples (axilla, groin, axilla-groin and nares swabs) containing either high or medium Ct values based upon the manual assay, were tested in quadruplicate. The BD MAX™️ assay was highly reproducible.

| **Sample No.** | ***C. auris* BD MAX^™^ Ct** | **Mean Ct ±**  **SD** | **%CV** |
| --- | --- | --- | --- |
| High Ct-1 | 29.0 | 28.9 **±**  0.2 | 0.8 |
| High Ct-2 | 28.9 |  |  |
| High Ct-3 | 29.0 |  |  |
| High Ct-4 | 28.5 |  |  |
| Medium Ct-1 | 24.3 | 25.3 **±**  0.7 | 3.0 |
| Medium Ct-2 | 25.3 |  |  |
| Medium Ct-3 | 26.1 |  |  |
| Medium Ct-4 | 25.5 |  |  |

**Supplementary Table 2. Specificity of BD MAX™️ real-time PCR assay.** Two concentrations of different clades of *C. auris*, and closely-and distantly-related *Candida* species were tested. The assay was highly specific for all *C. auris* clades with no cross-reactivity with other *Candida* species. The Ct values for all four clades for a given dilution were similar, further confirming the potential utility of the assay for all *C. auris* clades.

| Fungus (accession no.*) | CFU/PCR Reaction | Manual Ct | BD MAX™ Ct |
| --- | --- | --- | --- |
| *C. auris* South Asia Clade (M5658) | 1 x 10^4^ | 21.51 | 23.3 |
|  | 1 x 10^1^ | 33.35 | 34.4 |
| *C. auris* South Africa Clade (M5948) | 1 x 10^4^ | 22.45 | 23.5 |
|  | 1 x 10^1^ | 37.02 | 34.8 |
| *C. auris* South America Clade (M5952) | 1 x 10^4^ | 21.17 | 21.3 |
|  | 1 x 10^1^ | 32.26 | 33.1 |
| *C. auris* East Asia Clade (M5956) | 1 x 10^4^ | 17.84 | 19.5 |
|  | 1 x 10^1^ | 31.71 | 34.2 |
| *C. haemulonii* (M8541) | 1 x 10^4^ | Undet | -1 |
|  | 1 x 10^1^ | Undet | -1 |
| *C. duobushaemulonii* (M3051) | 1 x 10^4^ | Undet | -1 |
|  | 1 x 10^1^ | Undet | -1 |
| *C. albicans* (M3223) | 1 x 10^4^ | Undet | -1 |
|  | 1 x 10^1^ | Undet | -1 |
| *C. glabrata* (M8984) | 1 x 10^4^ | Undet | -1 |
|  | 1 x 10^1^ | Undet | -1 |

* NYSDOH Mycology Laboratory Culture Collection Repository

**Supplementary Table 3 A - Inter-assay Reproducibility of BD MAX™️ real-time PCR assay.**

Eight pools of surveillance samples (axilla, groin, axilla-groin and nares swabs) with varying Ct values based upon the manual assay, were tested by the BD MAX^TM^ assay on three different days. The consistent results revealed high inter-assay reproducibility.

| **Calculated CFU/PCR Reaction** | **BD MAX^™^ *C. auris* target** | | | | | |
| --- | --- | --- | --- | --- | --- | --- |
|  | **Run 1**  **Ct** | **Run 2**  **Ct** | **Run 3**  **Ct** | **Mean Ct** | **Ct SD** | **%CV** |
| 96294 (~100000) | 20.5 | 20.1 | 20.5 | 20.4 | 0.2 | 1.1 |
| 3094 | 25.4 | 25.6 | 26.0 | 25.7 | 0.3 | 1.2 |
| 284 | 29.3 | 30.0 | 30.7 | 30.0 | 0.7 | 2.3 |
| 46 | 31.4 | 32.0 | 32.0 | 31.8 | 0.3 | 1.1 |
| 29 | 33.5 | 36.3 | 34.6 | 34.8 | 1.4 | 4.1 |
| 10 | 39.4 | 36.7 | 37.2 | 37.8 | 1.4 | 3.8 |
| 1 | 37.6 | 35.3 | 39.7 | 37.6 | 2.3 | 6.0 |
| 0.4 (~0.5) | -1.0 | -1.0 | -1.0 | -1.0 | 0.0 | NA |

**Supplementary Table 3 B - Intra-assay Reproducibility of BD MAX™️ real-time PCR assay.** Two pools of surveillance samples were tested in quadruplicate on the same day. The consistent results revealed high intra-assay reproducibility.

| **Sample Type** | **BD MAX^™^ *C. auris* target** | | | |
| --- | --- | --- | --- | --- |
|  | **Ct** | **Mean Ct** | **Ct SD** | **% CV** |
| Pool 1  (axilla-groin) | 27.1 | 27.0 | 0.5 | 1.8 |
|  | 27.2 |  |  |  |
|  | 27.4 |  |  |  |
|  | 26.3 |  |  |  |
| Pool 2  (nares) | 22.8 | 22.8 | 0.4 | 1.7 |
|  | 22.2 |  |  |  |
|  | 22.9 |  |  |  |
|  | 23.1 |  |  |  |
